# Calumenin contributes to epithelial-mesenchymal transition and predicts poor survival in glioma

**DOI:** 10.1101/2020.07.05.188318

**Authors:** Ying Yang, Jin Wang, Shihai Xu, Fei Shi, Aijun Shan

**Affiliations:** Department of Pediatrics, Futian Women and Children Health Institute, Shenzhen 518045, China; Department of Emergency, Shenzhen People’s Hospital (The Second Clinical Medical College◻Jinan University; The First Affiliated Hospital, Southern University of Science and Technology), Shenzhen 518020, China

**Keywords:** Calumenin, CALU, glioma, epithelial-mesenchymal transition, EMT

## Abstract

Calumenin (CALU) has been reported to be associated with invasiveness and metastasis in some malignancies. However, in glioma, the role of CALU remains unclear. In the current study, we aimed to unveil its role in glioma based on transcriptome level. Clinical and transcriptome data of 998 glioma patients, including 301 from CGGA mRNA microarray dataset and 697 from TCGA RNA sequencing dataset, were downloaded and analyzed. R language was used to perform statistical analyses and generate figures. In glioma, CALU expression seemed to be positively associated with WHO grade system, and was enriched in IDH wildtype, mesenchymal and classical subtype. Genes that tightly correlated with CALU were screened and annotated with Gene Ontology, and it turned out that, these genes were highly enriched in cell/biological adhesion, response to wounding, and extracellular matrix/structure organization, all of which were strongly correlated with the epithelial-mesenchymal transition (EMT) phenotype. Subsequent GSEA analysis further validated the profound involvement of CALU in EMT. To get further understanding of the association between EMT and CALU, GSVA analysis was performed to identify the EMT signaling pathways that CALU might involve. CALU expression was found to be positively correlated with TGFβ, PI3K/AKT, and hypoxia pathway. Furthermore, Pearson correlation indicated that CALU played synergistically with EMT key markers, including N-cadherin, vimentin, snail, slug and TWIST1, in both CGGA and TCGA dataset. Kaplan-Meier curves and Cox regression analyses showed that higher CALU predicted a worse survival for patients, and the prognostic value was independent of WHO grade and age. In conclusion, CALU was correlated with more malignant phenotypes in glioma. Moreover, CALU seemed to serve as a pro-EMT molecular target and could contribute to predict prognosis independently for glioma patients.

## 1. Introduction

In central nervous system, glioma is the most prevalent and fatal primary cancer in adults^[1]^. Despite a substantial body of improvements in therapy, the prognosis for most glioma patients is still dismal. Particularly for patients who suffered from higher grade glioma (WHO grade IV, glioblastoma, GBM), which is the most malignant and lethal type, the median survival remains less than 15 months^[2, 3]^. There is a growing recognition that epithelial-mesenchymal transition (EMT) plays a key role in mediating tumorigenesis, stemness, invasiveness, resistance to radiochemotherapy, and early recurrence in glioma^[4–7]^. It is therefore imperative to identify novel EMT-related molecules for potential glioma diagnosis and intervention.

Calumenin (CALU) has been widely reported in a range of malignancies including head and neck cancer^[8]^, endometrial cancer^[9]^, colon^[10]^ and colorectal cancer^[11]^, lung cancer^[10, 12]^, melanoma^[13]^, hepatocellular and pancreatic carcinoma^[14]^, and breast cancer^[15]^. CALU, a calcium-binding protein localized in the endoplasmic reticulum (ER), is mainly involved in such ER functions as protein folding and sorting. Besides, CALU has recently been shown to influence cell mobility, migration, invasion, and metastasis during particular events, such as tumorigenesis, wound healing, immune response and coagulation^[16–21]^. Several studies have explored the relationship between CALU expression and survival and yielded relatively consistent results. In most types of cancer, a higher level of CALU in lesions indicated a more malignant phenotype and a shorter survival for patients.

However, the expression patterns and biological functions of CALU in gliomas have rarely been described. Only one study presented by Sreekanthreddy et al.^[22]^, investigated the prognostic potential of serum CALU in GBM, which accounted for about 40% of pan-glioma. Here, we analyzed clinical and transcriptome data of 998 patients, aiming at exploring the role of CALU in gliomas.

## 2. Materials and Methods

### 2.1. Sample and data collection

From Chinese Glioma Genome Atlas website (CGGA, http://www.cgga.org.cn/), we selected 301 glioma samples with mRNA microarray data. From The Cancer Genome Atlas website (TCGA, http://cancergenome.nih.gov/), we obtained 697 glioma patients with RNA-sequencing data. Clinical data, including WHO grade, IDH mutation status, molecular subtype, and prognosis, were also available. Thus, a total of 998 samples were included in the present study. Baseline characteristics of glioma samples in both datasets were summarized in Table S1. In CGGA_301 dataset, microarray data, which had already been normalized and centered (using GeneSpring GX 11.0 platform) by data provider, were directly utilized. While in TCGA_697 dataset, RNAseq data (RSEM normalized, level 3) were log2 transformed before data analysis.

### 2.2. Statistical analysis

Statistical analyses were primarily performed with R language (version 3.6.2). A set of R packages, such as ggplot2, pROC^[23]^, pheatmap, corrgram, circlize^[24]^, and gsva, were used to handle corresponding calculations and to produce figures. Cox proportional hazard regression analyses were performed with coxph function of survival package. Gene ontology analysis (GO) of CALU-correlated genes was implemented based on DAVID^[25]^ website (version 6.8, https://david-d.ncifcrf.gov/). For Gene Set Enrichment Analysis (GSEA)^[26]^ and Gene Set Variation Analysis (GSVA)^[27]^, a series of gene sets were obtained from the GSEA network (http://software.broadinstitute.org/). A p-value less than 0.05 was considered to be statistically significant. Two-sided significance tests were adopted throughout.

## 3. Results

### 3.1. CALU was significantly upregulated in GBM, IDH wildtype, mesenchymal, and classical subtype

According to the WHO grade system, CALU expression was analyzed in both CGGA and TCGA dataset, and the results congruently showed a significantly positive correlation between WHO grade and CALU expression (Figs. 1A and C). Moreover, when IDH mutation status was defined as a subclassifier, we observed that IDH wildtype GBM exhibited the highest expression pattern of CALU in both CGGA and TCGA dataset. Besides, CALU expression in IDH mutant glioma seemed to be universally lower than that in IDH wildtype, across different WHO grades, except for lower grade glioma (LGG) in CGGA, which exhibited apparent trends although not significant (Figs. 1B and E). Subsequently, the distribution of CALU expression among different molecular subtypes was investigated. As shown in Figs. 1C and F, CALU was significantly upregulated in classical and mesenchymal subtype compared to neural and proneural subtype. These findings indicated that higher CALU expression was usually accompanied by higher malignancy potential of glioma.

**Fig. 1.**
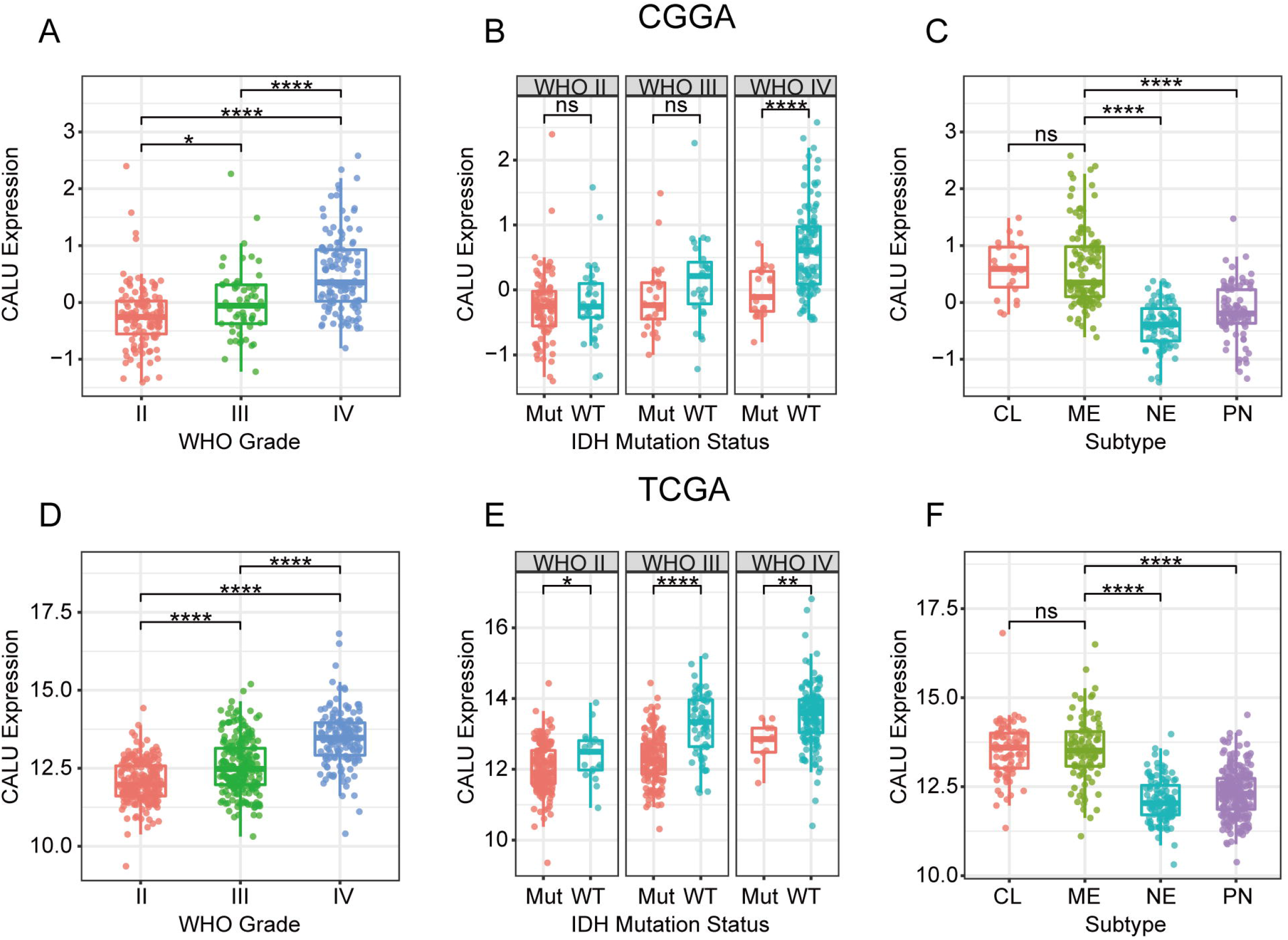
CALU expression in CGGA and TCGA dataset according to WHO grade (A, D), IDH mutation status (B, E), TCGA molecular subtype (C, F). * indicates p value < 0.05, * * indicates p value < 0.01, * * * indicates p value < 0.001, * * * * indicates p value < 0.0001.

### 3.2. CALU-related biological process

We performed Pearson correlation to calculate the correlation coefficient between CALU and every gene. Genes that strongly correlated with CALU were screened out with Pearson |r| > 0.6 in each dataset. In total, 621 genes in CGGA chort and 965 in TCGA cohort were eligible. To ensure the accuracy of the analysis, we subsequently identified 203 genes that overlapped between two independent cohorts, all of which were positively correlated with CALU (Table S2). Based on these genes, GO analysis revealed that, genes that significantly correlated with CALU were highly enriched in a set of biological processes that correlated with EMT, including cell/biological adhesion, response to wounding, extracellular matrix/structure organization, collagen fibril organization, and collagen biosynthetic process (Figs. 2A and B). Moreover, the association between CALU expression and EMT was revealed by GSEA analysis. CALU expression was found to be positively associated with the gene set of HALLMARK_EPITHELIAL_MESENCHYMAL_TRANSITION in both CGGA dataset (NES = 1.897, FDR = 0.035) and TCGA dataset (NES = 1.818, FDR = 0.075) (Figs. 2C and D). These findings suggested that CALU might be particularly involved in EMT process during glioma progression.

**Fig. 2.**
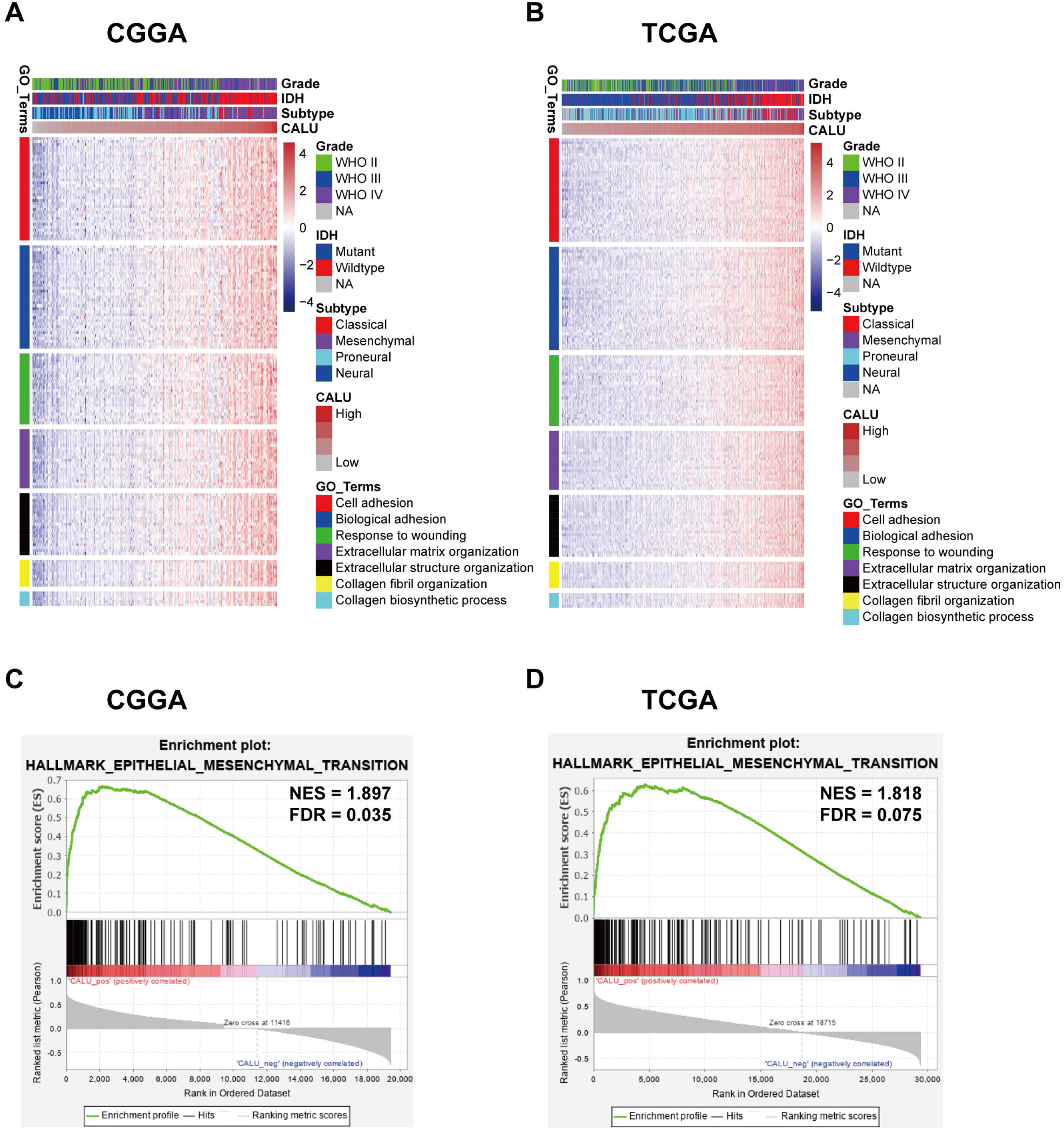
Functional enrichment of CALU in glioma. Gene Ontology analysis (A, B) and Gene set enrichment analysis (C, D).

### 3.3. CALU-related EMT signaling pathways

To get further understanding of the association between CALU and EMT, Seven gene sets, representing distinct EMT signaling pathways^[28]^ were obtained from GSEA network (Table S3). Through cluster analyses, we identified 3 EMT signaling pathways (TGF-β, PI3K/AKT, and hypoxia), which might be strongly correlated with CALU (Figs. 3A and B). Moreover, seven gene sets were transformed into seven metagenes with GSVA analysis, which were subsequently put into Pearson correlation together with CALU. According to Pearson r among seven metagenes and CALU, Corrgrams were plotted to assess their interrelationships. CALU was found to be positively correlated with TGF-β, PI3K/AKT, and hypoxia, in line with what we observed in clusters. While only a very weak correlation was revealed between CALU expression and four other pathways (WNT, MAPK, NOTCH, and HEDGEHOG), which might be ascribed to signal noise (Figs. 3C and D).

**Fig. 3.**
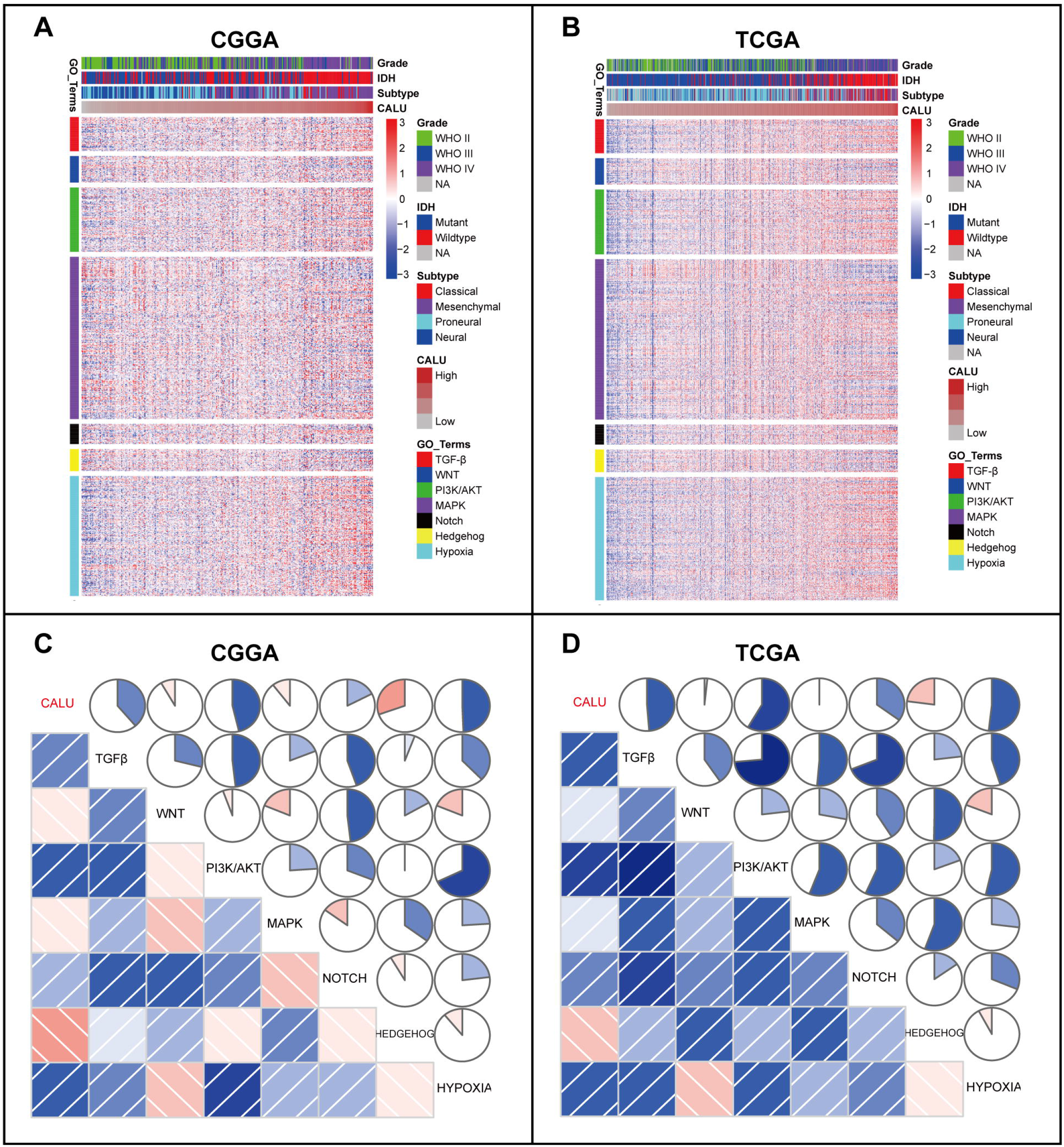
Cluster (A, B) and GSVA (C, D) of CALU-related EMT signaling pathways in glioma.

### 3.4. CALU was synergistic with EMT key markers

Assuming that CALU played a vital role in regulating glioma EMT, we investigated the association between CALU and EMT markers, including N-cadherin, E-cadherin, snail, slug, and vimentin. Pearson correlation tests were performed with CALU and the above five EMT markers in both CGGA and TCGA. Circos plots were derived from Pearson r-values between CALU and five markers. As shown in Figs. 4A and B, CALU expression showed high agreement with N-cadherin, snail, slug, and vimentin. In contrast, a weak relationship between CALU and E-cadherin was found in Circos plots, which could be defined as a noise. Heretofore, some other members, including TWIST1/2, β-catenin, and ZEB1/2, have been reported as key markers in EMT^[29]^. Thus, we additionally put them into analysis together with CALU. CALU expression was tightly associated with TWIST1 in both CGGA and TCGA dataset (Figs. 4C and D).

**Fig. 4.**
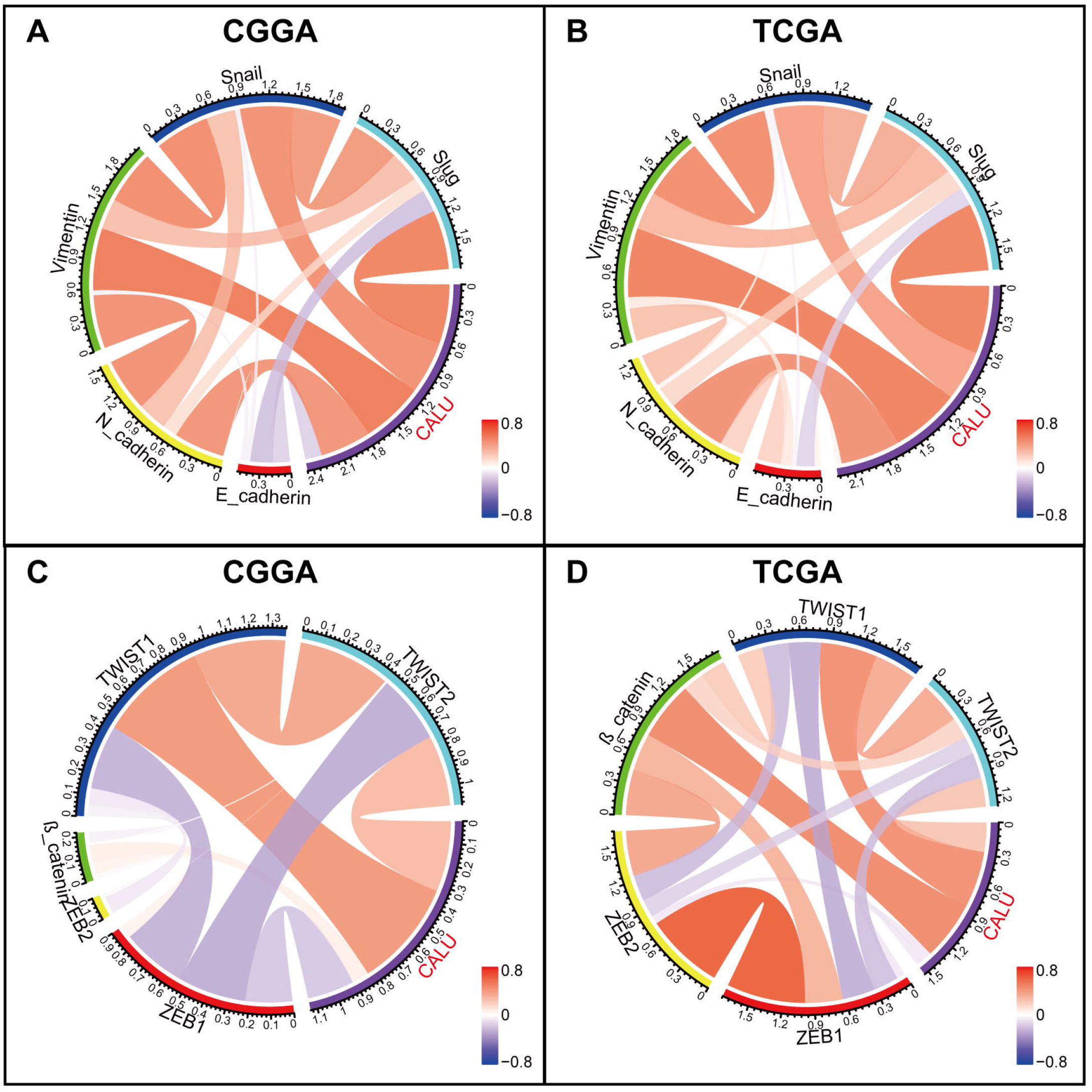
Correlation of CALU and EMT key biomarkers.

### 3.5. Higher CALU was related to a worse prognosis

To evaluate the prognostic value of CALU in glioma, Kaplan-Meier (KM) survival curves were plotted. In pan-glioma samples, when comparing the two groups defined by median CALU expression, we observed that higher CALU expression predicted a significantly shorter survival, as shown in Figs. 5A and D. Moreover, glioma patients were further divided into LGG and GBM subgroup. In both subgroups, patients with higher CALU exhibited universally worse survival than those with relatively lower CALU (Figs. 5B, C, E and F), except for TCGA GBM, which also showed an apparent trend. To identify the independent effect of CALU on glioma prognosis, Cox regression analyses were performed with covariates including CALU expression, age and WHO grade. Multivariate analyses revealed that CALU expression was a significant prognosticator independent of age and WHO grade in both CGGA and TCGA (Table 1).

**Fig. 5.**
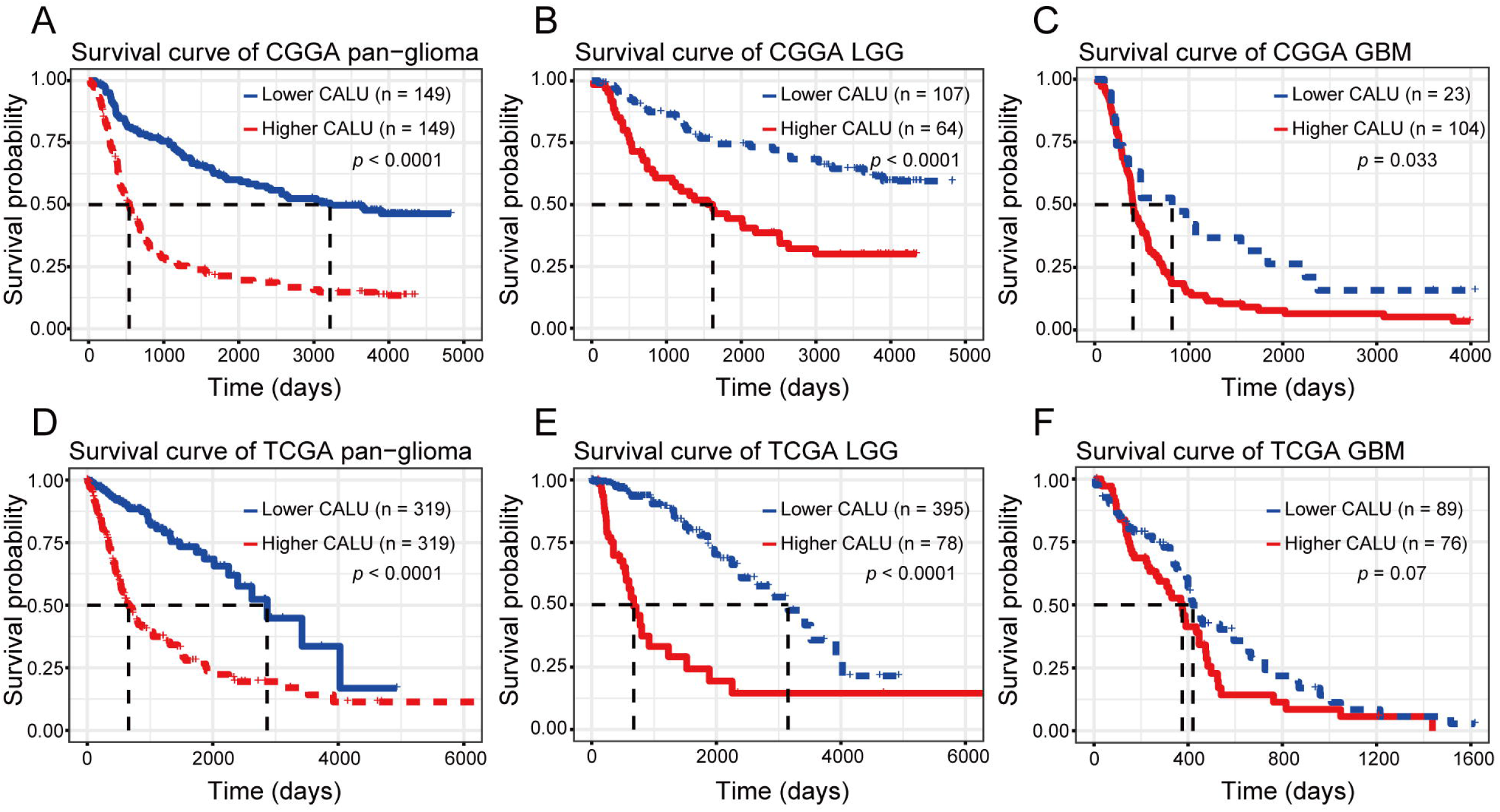
Survival analysis for CALU in pan-glioma (A, D), LGG (B, E) and GBM (C, F).

**TABLE 1.** Cox regression analysis of overall survival in glioma

| Covariates | CGGA_301 |  |  |  | TCGA |  |  |  |
| --- | --- | --- | --- | --- | --- | --- | --- | --- |
|  | Univariate |  | Multivariate |  | Univariate |  | Multivariate |  |
|  | HR(95% CI) | *P* | HR(95% CI) | *P* | HR(95% CI) | *P* | HR(95% CI) | *P* |
| Age | 1.041<br>(1.027-1.055) | 0.000 | 1.017<br>(1.004-1.031) | 0.010 | 1.075<br>(1.062-1.087) | 0.000 | 1.046<br>(1.032-1.060) | 0.000 |
| Grade | 2.670<br>(2.221-3.210) | 0.000 | 2.292<br>(1.867-2.814) | 0.000 | 5.057<br>(3.915-6.532) | 0.000 | 3.033<br>(2.273-4.047) | 0.000 |
| CALU | 2.123<br>(1.746-2.581) | 0.000 | 1.300<br>(1.033-1.636) | 0.025 | 2.159<br>(1.892-2.463) | 0.000 | 1.295<br>(1.094-1.534) | 0.003 |

## 4. Discussion

We explored CALU expression at transcriptional level via a cohort of 998 glioma samples, and demonstrated that CALU expression was positively correlated with WHO grades. In addition, upregulation of CALU was usually paralleled with a more malignant and aggressive phenotype, such as IDH wildtype, classical subtype and mesenchymal subtype. Survival analyses revealed that higher CALU was related to a worse prognosis, independent of age and WHO grade. These results concordantly indicated that CALU might contribute to malignant progression of glioma, which were in line with the results from a previous GBM study^[22]^. Thus, unveiling the regulative mechanism of CALU may facilitate to develop a novel gene for potential glioma diagnosis and treatment.

Calumenin is one of the members of CREC protein family. This molecule family mainly consists of Cab45, Reticulocalbin1, ERC-55 and Calumenin, and is characterized by multiple EF-hand motifs with low affinity of Ca^2+^-binding^[30]^. Under normal physiological conditions, CALU primarily participates in regulating Ca^2+^-dependent protein folding, sorting and maturation in the ER^[31]^, Ca^2+^ homeostasis^[32, 33]^, and muscle contraction/relaxation^[34]^. While in tumor microenvironment, CALU was reported to play a critical role in promoting a series of malignant phenotypes including cancer cell survival^[21]^, filopodia formation and cell migration^[20]^, invasiveness^[12]^, metastasis^[15, 35]^, cancer development^[10]^, and resistance to chemotherapy^[13]^. So far very little is known about the biological function of CALU in glioma. In the current study, GO analysis was performed to elucidate the biological function of CALU in glioma, and it revealed that, CALU showed highly association with multiple EMT-related biological processes, including cell adhesion, biological adhesion, extracellular matrix/structure organization, collagen fibril organization, and collagen biosynthetic process. GSEA in both CGGA and TCGA further exhibited a remarkable relationship between CALU and EMT. EMT have been extensively reported to act as a critical mechanism not only in invasiveness but also in early recurrence and resistance to therapy in glioma^[5, 6, 36]^. These findings suggested that CALU might facilitate the malignant progression of glioma primarily via modulating EMT process, which has not yet been reported previously. Despite no report with regard to the pro-EMT effect of CALU, the other two members (Cab45^[37]^ and EFHD2^[38]^) from the same protein family have been described in EMT regulation, which indirectly supported the potential role of CALU in glioma EMT.

We then chose a panel of EMT pathways and markers, and examined their interrelationships with CALU. CALU was revealed to be highly associated with TGFβ, PI3K/AKT, as well as hypoxia pathway, indicating that CALU might regulate glioma EMT through these signaling pathways. Furthermore, most of the EMT biomarkers showed robust correlation with CALU, suggesting a synergistic effect among CALU and these members during EMT process. These findings further validated the potential role of CALU during EMT process in glioma.

## Conclusions

In conclusion, CALU was upregulated in more malignant gliomas, and predicted much worse prognosis. Furthermore, CALU seemed to be mainly involved in EMT process of glioma, potentially through modulating TGFβ, PI3K/AKT, and hypoxia pathway. However, the conclusions still need to be validated in further experimental studies.

## Acknowledgments

We appreciate the generosity of CGGA project and TCGA project for sharing data. This work was supported by Medical Scientific Research Foundation of Shenzhen Health Commission (szfz2018022) and Science and Technology Innovation Foundation of Shenzhen (JCYJ20190806150005453).

